# MCIBox: A Toolkit for Single-molecule Multi-way Chromatin Interaction Visualization and Micro-Domains Identification

**DOI:** 10.1101/2022.04.28.489957

**Authors:** Simon Zhongyuan Tian, Guoliang Li, Duo Ning, Kai Jing, Yewen Xu, Yang Yang, Melissa J. Fullwood, Pengfei Yin, Guangyu Huang, Dariusz Plewczynski, Wenxin Wang, Jixian Zhai, Ziying Wang, Ziwei Dai, Yuxin Lin, Wei Chen, Meizhen Zheng

## Abstract

The emerging ligation-free three-dimensional (3D) genome mapping technologies can identify multiplex chromatin interactions with single-molecule precision. These technologies offer new insight into high-dimensional chromatin organization and gene regulation, but also introduce new challenges in data visualization and analysis. To overcome these challenges, we developed MCIBox, a toolkit for Multi-way Chromatin Interaction (MCI) analysis, including a visualization tool and a platform for identifying micro-domains with clustered single-molecule chromatin complexes. MCIBox is based on various clustering algorithms integrated with dimensionality reduction methods that can display multiplex chromatin interactions at single-molecule level, allowing users to explore chromatin extrusion patterns and super-enhancers regulation modes in transcription, and to identify single-molecule chromatin complexes that are clustered into micro-domains. Furthermore, MCIBox incorporates a two-dimensional kernel density estimation algorithm to identify micro-domains boundaries automatically. These micro-domains were stratified with distinctive signatures of transcription activity and contained different cell cycle associated genes. MCIBox could potentially distinguish the specificity of single-molecule chromatin interaction patterns in various phases of a cell cycle or cell types.

## INTRODUCTION

Eukaryotic genomes are organized into a three-dimensional (3D) multiscale structure within the nucleus, and this structure is often associated with gene regulation (Dekker, 2008). The proximity ligation-based 3D genome mapping technologies, such as Hi-C and ChIA-PET, have enabled global mapping of chromatin interactions and characterization of nuclear genome organization in multiple scales, ranging from loops, Topologically Associated Domains (TADs), to A/B compartments (Fullwood et al., 2009; Li et al., 2012; Lieberman-Aiden et al., 2009; Rao et al., 2014; Tang et al., 2015). State-of-the-art visual tools such as Juicebox, HiGlass, 3D Genome Browser, WashU Epigenome Browser, BASIC Browser, and Nucleome Browser (http://vis.nucleome.org/entry/) have been developed for users to explore 3D genome organization and long-range chromatin interactions conveniently (Durand et al., 2016; Kerpedjiev et al., 2018; Lee et al., 2020; Wang et al., 2018; Zhou et al., 2013) (Zhu et al., 2022). Thereinto, BASIC Browser displays RNA polymerase II (RNAPII) mediated promoter-enhancer contacts as complex chromatin interactions, named as RAIDs (RNAPII-mediated chromatin interaction domains), which offers the genomics evidence for the proposed model of “transcription factories” from RNAPII foci observed via imaging data. These tools have provided deep insight into the principle of gene regulation.

However, the aforementioned experimental methods and visualization tools can only capture pairwise interactions. Moreover, these methods present accumulated results from bulk cell populations, thus they cannot precisely locate the interacting loci with high heterogeneity in single nuclei. Therefore, it remains unclear whether these multiplex chromatin interactions happen in individual nuclei or composed by different nuclei and finally showing as accumulated results. Thus, these methods cannot reveal the detailed nature of chromatin contacts for answering questions regarding the modes for TADs in individual nuclei or the composed pattern of super-enhancers associated with key genes for cell destination.

In order to overcome these limitations, a few ligation-free technologies have been developed. GAM, by sequencing a collection of thin cryosectioned nuclear profiles for genome architecture mapping (Beagrie et al., 2017), revealed three-way contacts among super-enhancers for specific gene regulation. SPRITE, a split-pool recognition of chromatin interactions by tag extension (Quinodoz et al., 2018), showed “active hub” around nuclear speckles enriched with RNAPII activity and “inactive hub” around nucleolus enriched rDNA corresponding to low transcribed activity. ChIA-Drop, which is the microfluidic-based and barcode-linked sequencing for chromatin-interaction analysis (Zheng et al., 2019), showed a strong directionality bias towards the gene body for supporting one-sided extrusion model of transcription. These methods have enabled users to capture multiple interacting genomic loci simultaneously.

ChIA-Drop, SPRITE and GAM approaches have demonstrated the multi-way chromatin interactions involved in the 3D genome architecture and gene regulation and comprehensively mapped chromatin contacts at a previously unappreciated level (Kempfer and Pombo, 2020). They also provided evidence that the single-molecule chromatin complexes with high heterogeneity and also clustered together as microscale domains via certain similarity, contributing to the chromatin structure such as TADs and RAIDs as accumulated results (Zheng et al., 2019). The emerging evidence indicates that chromatin microscale domains that are organized into TADs exhibit higher tendency of cell-type-specific 3D genome structure than low-resolution TADs (Phillips-Cremins et al., 2013). These ligation-free methods can elucidate principles of genome folding at microscale by clustering their similarity and can demultiplex the averaged interacting loci precisely into single nuclei multi-way interactions. In this study, for the microscale domain constructed into TADs were called microTADs, and that constructed into RAIDs associated with transcription factories were called microTFs, respectively, for further investigation.

So far, these novel exciting findings were identified with computational methods and integrative analysis. However, the visualization tools for real-time detailed profiling of single-molecule chromatin complexes and the characterization of micro-domains are currently lacking. Here we report MCIBox, a new toolkit for profiling multi-way chromatin interactions and visualizing micro-domains clustered from single-molecule chromatin interaction complexes. These approaches include the hierarchical clustering algorithm on data matrix from GAM, SPRITE and ChIA-Drop datasets respectively, or various clustering algorithms on scatter plot matrix created with dimensionality reduction on that data matrix. Our MCIBox provides a convenient interface “MCI-view” for real-time exploration of chromatin topological structure and higher-order chromatin organization in multi-way chromatin interactions data, such as the chromatin extrusion patterns in gene transcription and chromatin organization, the working modes of super-enhancers in transcription regulation (Beagrie et al., 2017).

When investigating the ligation-free data of ChIA-Drop, GAM and SPRITE by MCI-view, we found that individual chromatin TADs contained several micro-domains. Thus, we added a program in MCIBox for micro-domain characterization, a Two-Dimensional Kernel Density Estimation (2D KDE) contour map based on “MCI-2kde”. By focusing on an RNAPII associated ChIA-Drop dataset of *Drosophila* S2 cells (Zheng et al., 2019), we applied MCI-2kde program to automatically determine the boundary of micro-domains, which are micro-Transcription factories (microTFs) mediated by RNAPII transcription factors. We identified 578 microTFs from the 126 of 476 RAIDs at length scales above 150 kilobase (kb) that determined by previously pairwise RNAPII ChIA-PET data (Zheng et al., 2019). These microTFs were stratified from various patterns with distinctive signatures of transcription activity and histone modification. Interestingly, different microTFs in a RAID contained specific genes in the different phases of a cell cycle, demonstrating that MCIBox can potentially distinguish single-molecule chromatin interactions with cell-cycle or cell-type specificity.

Taken together, MCIBox represents an invaluable tool for the study of multiple chromatin interactions and inaugurates a previously unappreciated view of 3D genome structure.

## RESULTS

### Overview of MCIBox

There are two main functions in MCIBox: a browser “MCI-view” for multiple chromatin interaction data visualization, and a framework for micro-domain boundary quantification using “MCI-2kde” program. MCI-view is for the visualization of single-molecule chromatin interaction complexes from the emerging ligation-free based approaches including ChIA-Drop, SPRITE and GAM (Figures 1). MCI-view can be applied to view new aspects of 3D genome topology and microscale chromatin structure, such as the chromatin fiber organization activity during transcription and regulation, the single-molecule chromatin complexes clustered micro-domains in TADs (called microTADs), or the micro-domains of transcription factories (TFs) (named microTFs). MCI-2kde is a two-dimensional kernel density estimation algorithm based unsupervised machine learning method for micro-domain definition automatically.

**Figure 1.**
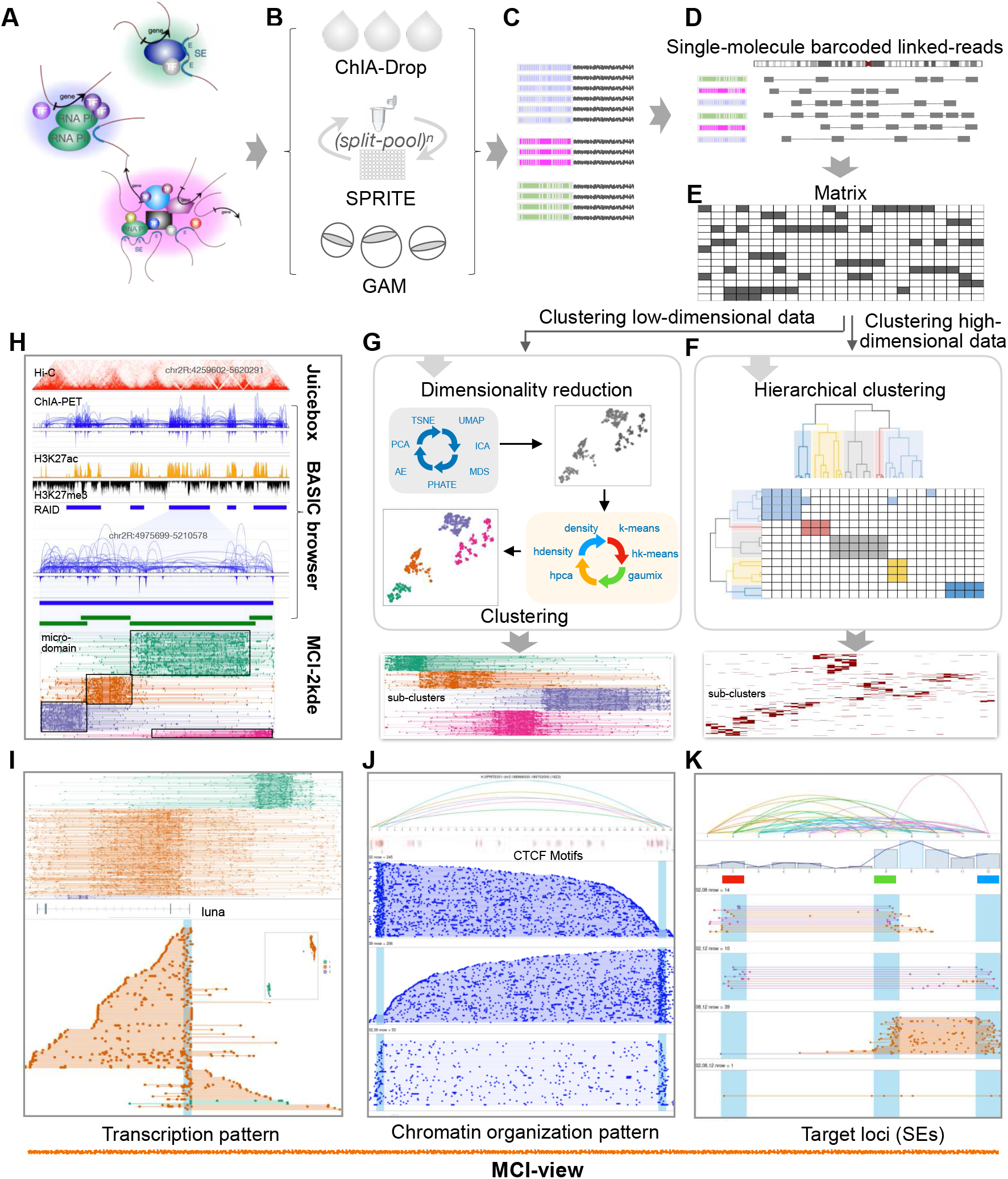
Overview of MCIBox for multi-way chromatin interaction analysis. (A-C) Schematic for generating single-molecule multi-way chromatin interaction datasets with different ligation-free based methods including ChIA-Drop, SPRITE, and GAM. (A) for chromatin complexes preparation, (B) for different methods, and (C) for sequencing reads that were grouped by barcodes. (D) A multiplex chromatin interaction complex is defined as the fragments with the same barcodes. (E) The chromatin complexes from the genomic region of interest were used as the input into MCIBox for matrix construction. Binned genomic regions are as columns along x-axis, and binned fragments from the complexes are arranged as rows along y-axis. (F) Subsequently, these binned fragments were arranged by similarity via hierarchical clustering upon high-dimensional data, which are ready for Cluster-view (a visualization function in MCIBox). (G) Using dimensionality reduction algorithms (UMAP by default), the matrix was converted to 2-dimensional (2D) scatter plot, among them the tethered closely dots were clustered together by default algorithms of hierarchical k-means (hk-means) for Fragment-view (a visualization function in MCIBox). (H) A screenshot showing micro-domain from multi-way contacts with MCI-view, along with 2D heatmap of Hi-C with Juicebox, and pairwise interaction loops and coverage from ChIA-PET with BASIC Browser. The blue bar indicates RNAPII associated interaction domain (RAID), and the green bar indicates micro-domain (microTADs or microTFs). (I-K) Screenshots showing “Transcription pattern”, “Chromatin organization pattern”, and “Target loci” for multi-way contacts with MCI-view respectively.

### MCI-view displays contact maps for single-molecule multi-way chromatin complexes

MCI-view is a key component of MCIBox, and provides a visualization system for multi-way chromatin interactions. Users can visualize their own experimental data by creating a formatted document that contains information about complexes in each line from the results of multiple chromatin contacts generated by multi-way contacts detected approaches (Figures 1A-D and S1; Methods). MCI-view reads the data of the genomic region of interest from the formatted input file for matrix reconstruction (Figure 1E), then integrates hierarchical clustering strategies to cluster the high-dimensional data directly, or performs dimensionality reduction methods to reduce the matrix data into low-dimensional data for further clustering when the single-molecule complexes data feature is too complicated. This approach was specifically developed for integrative genomics viewer for handling multi-way contact data (Figures 1F and 1G). MCI-view also can browse accumulated density results for 1D track similar to the coverage of ChIP-seq data, and can browse 2D profiles such as loop or domain annotations, side by side simultaneously (Figure 1H).

For single-molecule chromatin interactions, the interface of MCI-view supplies a function to display the data by specifying the interested single or multiple genomic coordinates, or the interested gene name, and an additional function to filter out the uninterested data. Currently MCI-view include 6 modules for display multi-way contact results (Figures S1): (i) Cluster-view performs binning of clustering tracks for browsing the clustered microscale domains of multiplex chromatin interaction complexes (Figures S1D, S1G and S1K), (ii) Fragment-view directly visualizes the original (unbinned) fragments for the multi-way contacts representation (Figures S1E, S1H and S1I), (iii) Target loci view presents the specific region similarly to DNA-FISH probe associated genomic loci, selected by clicking on the interested regions (Figures 1K and S1I), (iv) Chromatin organization pattern view is used for observing the multiple chromatin interactions at transcription factor binding motifs such as CTCF motifs (Figures 1J and S1J), (v) Transcription pattern view is used for discovering the multiple chromatin interactions profiling during the process of gene transcription in order to explore the extrusion process (Figures 1I and S1K), (vi) Transcription regulation pattern view can be applied for observation of the multi-way chromatin interactions at super-enhancers region (Beagrie et al., 2017), thereby allowing users to identify the composition of super-enhancers for gene regulation.

### MCl-view uncovers super-enhancers interacting in single-molecule level

To explore the contact detail of super-enhancers with the working model as a whole unit or by different composed enhancers to regulate gene promoter in individual nuclei (Figure 2A), we used MCI-view to display the interacting profiling of super-enhancers from mouse ESC GAM data (Figure 2B, same genomic region in *Figure 5a* of the GAM paper from Beagrie et al.) (Beagrie et al., 2017). The 2D heatmap shows the pairwise contacts from GAM results with square highlight of enhancer contact regions. Cluster-view shows the different composed enhancers in the multiple chromatin complexes, along with the composition of micro-topologically associated domains (microTADs) that are accumulated into the TAD (Figure 2B). In addition, Fragment-view displays detailed original fragments clustering profiling for the super-enhancers located in microTADs (Figure 2C). This particular genomic region with three enhancers showed that 53.7% of multiplex chromatin complexes involve one enhancer, 37% involve two enhancers, and 9.3% of them involve three enhancers. Thus, the composition of enhancers can be variable in the super-enhancer region for gene-specific regulation in individual cells (Figure 2D).

**Figure 2.**
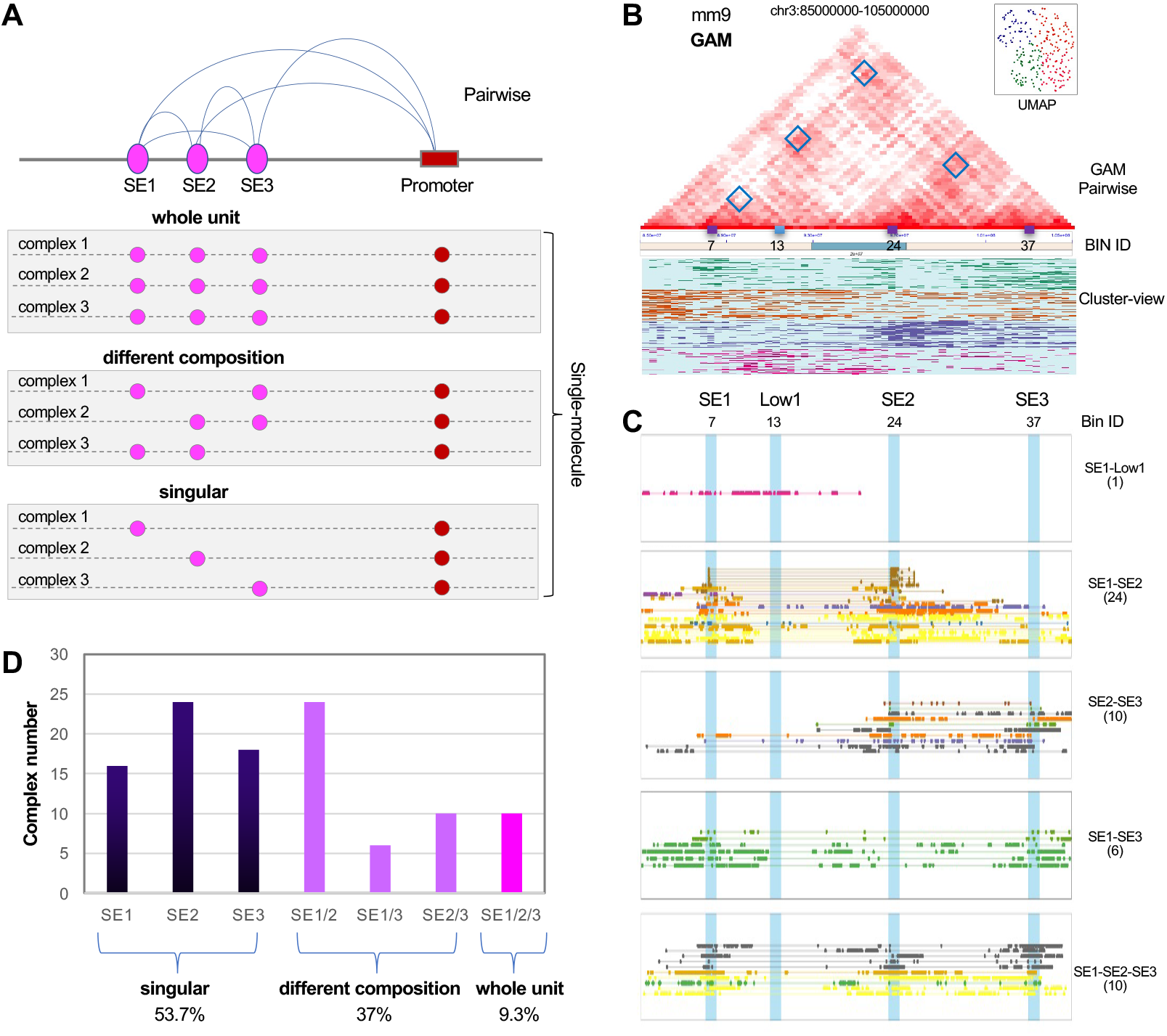
MCI-view displays super-enhancers working model. (A) Diagram of the potential working model for Super-enhancers (SEs) on gene regulation, “whole unit” working model presents all the enhancers in a super-enhancer region are tethered together to regulate target genes in different cells, “different composition” working model indicates different composed of enhancers co-regulate in different cells, “singular” working model shows only one enhancer regulates target genes in different cells. (B) MCI-view displays the pairwise contacts of the genomic region (chr3:85000000-105000000) from mouse GAM data, from which we could find the selected enhancer points annotated with square shown in the heatmap (*Figure 5a* from the GAM paper) (Beagrie et al., 2017). Besides this, the Cluster-view function in MCI-view shows that there are several micro-domains, which indicates different chromatin interaction clustering within this genomic region. Inset scatter plot presents the clustering algorithm. (C) “Transcription regulation pattern” in MCI-view to display the contact profiling of these super-enhancers regions. Number of “7, 13, 24 and 37” presents the Bin ID in this region and corresponding to “SE1, Low1, SE2 and SE3” in the GAM paper (Beagrie et al., 2017). SE1, SE2 and SE3 present super-enhancer 1, 2 and 3 loci. Low1 presents a control region not involved in the contacts among these super-enhancers. SE1-Low1 presents the interaction between super-enhancer SE1 and Low1 region, the number inside brackets shows the detected single-molecule complexes, so and forth. (D) Histogram shows the percentage of chromatin complexes for composed enhancers among SE1, SE2 and SE3.

### MCI-view directly visualizes asymmetric loop-extrusion process

Chromosomes are organized as chromatin loops to promote the interactions between enhancer-promoter, allowing for the long-range regulation of gene expression. Loops were hypothesized to form by “loop extrusion” and visualized by real-time imaging in vitro (Ganji et al., 2018). Our RNAPII ChIA-drop method captured multiplex interactions on *Drosophila* S2 cell line data presenting the asymmetric chromatin extrusion during gene transcription genome-wide. Here we applied human GM12878 SPRITE data to explore and display the loop extrusion associated with CTCF, which has not been analyzed in SPRITE paper since no published tools for visualization at that time.

The previously published SPRITE dataset (Quinodoz et al., 2018) was converted to the formatted document for MCI-view visualization. As shown in Figure S2A, the accumulated 2D heatmap shows pairwise contacts at the selected genomic region, while the 1D coverage shows variable density at this region corresponding to the higher-order structure of a domain on 2D heatmap. The key is that the Cluster-view module of MCI-view unfolds the layers for the detailed composition of multiple chromatin complexes in this region that were clustered into 9 micro-domains.

Fragment-view directly displays the original fragments information of these clustered chromatin complexes. Due to the MCI-view function for selecting the anchor of interests, we can clearly visualize the multi-way contacts by Cluster-view. In the case of this targeted region, the view agrees well with the previously published results (in “*Figure S2D*” of the SPRITE paper) (Quinodoz et al., 2018) and can further observe the original contact loci of the chromatin complexes by Fragment-view in MCI-view.

Moreover, we can observe the chromatin extrusion pattern from the multi-way contacts associated CTCF binding motifs via the module of “chromatin organization pattern view” in MCI-view (Figures S2B-D and Figure 3). In this genomic region (Figures 3A-D), there are 908 chromatin complexes that contained at least one CTCF motifs at left or right sides, and there are 461 of them that covered the left-side CTCF motifs, 364 of them that covered the right-side CTCF motif, only 83 (9%) of them covered both sides. These results indicate that CTCF-associated chromatin complexes exhibit one-sided DNA loop extrusion profiling, where CTCF complexes stop and tether at DNA loci with CTCF binding motifs, then reel the other side of DNA to close gradually and stop until meet another CTCF binding site with “convergent CTCF motifs”.

**Figure 3.**
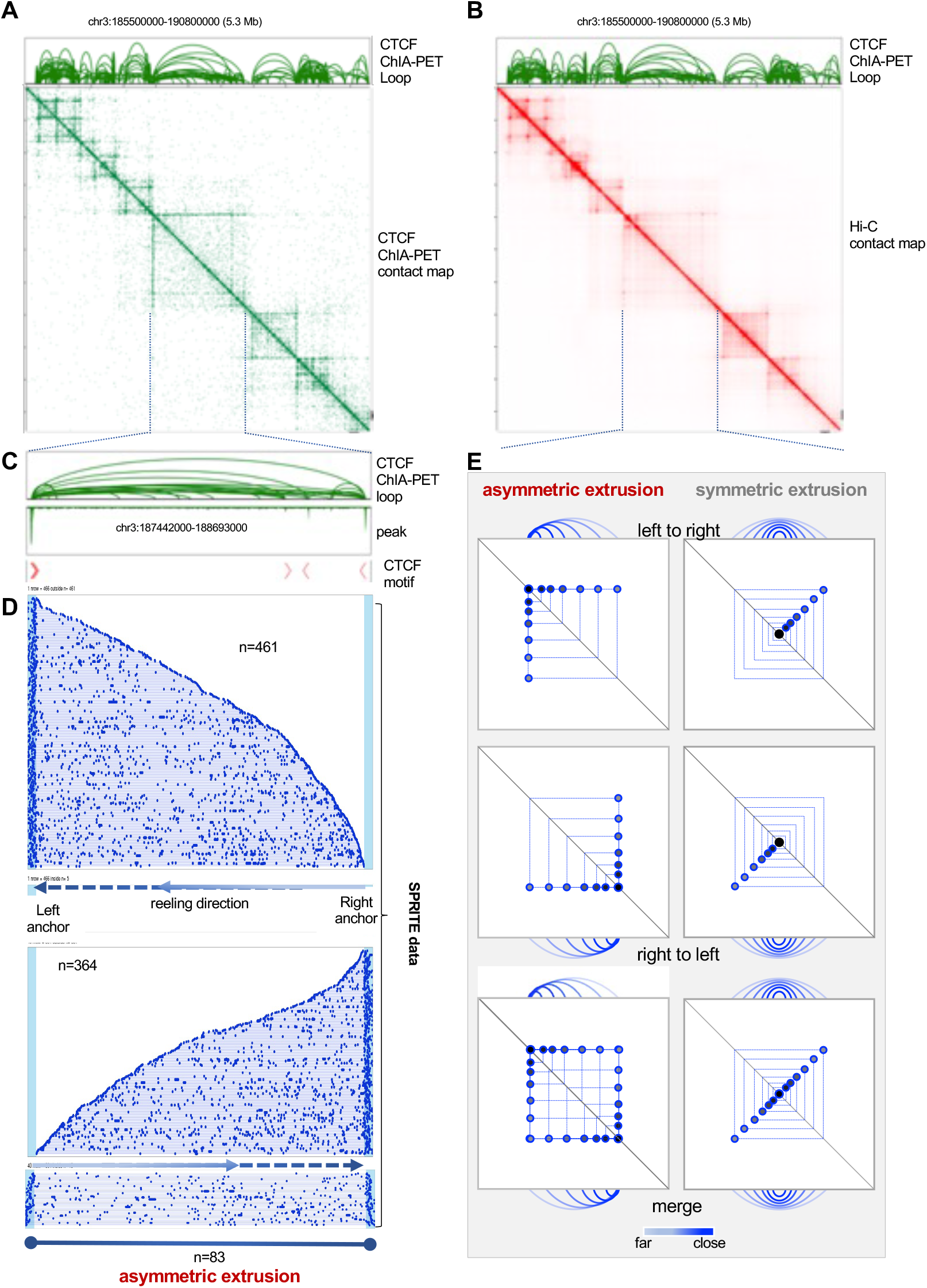
MCI-view displays chromatin loops extrusion. (A-B) BASIC Browser displays the pairwise contact loops of the genomic region (chr3:185500000-190800000) from CTCF ChIA-PET data on human GM12878 cells, along with 2D heatmap from CTCF ChIA-PET data (green) and Hi-C data (red). (C) BASIC Browser displays the pairwise contact loops and binding peaks of the zoomed-in genomic region (chr3:187442000-188693000) from CTCF ChIA-PET data on human GM12878 cells. Arrow with red color indicates CTCF binding motifs. (D) MCI-view displays the single-molecule chromatin complexes of the genomic region (chr3:187442000-188693000) from SPRITE data on human GM12878 cells. MCI-view displays SPRITE multiplex chromatin complexes covered the left anchor contained CTCF motifs with cyan color, multiplex chromatin complexes were shown as Fragment-view aligned by complexes span length in the local region, followed by that of right anchor and both anchors. Line with arrows present chromatin loops reeling direction, line with solid circle indicates chromatin loops stop reeling, the number (n) of chromatin complexes overlapped with CTCF motifs is shown. (E) Diagram of asymmetric and symmetric chromatin loops extrusion is shown. Arch diagram showing positions of one-sided or two-sided CTCF complexes translocating along chromatin DNA (grey) and progressively growing loops at different times indicated by color from blue (close) to transparent blue (far). The drawing 2D contacts corresponding the asymmetric and symmetric loops extrusion is shown under the arch diagram. Chromatin loops translocating along chromatin DNA from left to right (Top), right to left (Middle), merge of both (Bottom), are shown.

Our observation of loop extraction activity in MCI-view is consistent with the findings of real-time imaging of asymmetric DNA loop extrusion that condensin anchors onto DNA and reels it in from only one side (Ganji et al., 2018), and the findings of live-cell imaging of loops covered both sides are rare (Gabriele et al., 2022). Additionally, RNAPII ChIA-Drop results provide a promoter-centred extrusion model during gene transcription (Figure S2E) (Zheng et al., 2019). We note that the putative higher-order structure domain of 2D heatmap from Hi-C data supports the notion that chromatin loops extrude asymmetrically. Otherwise, if they extrude symmetrically, they would exhibit a secondary diagonal overlaid on the main diagonal as a “+” shaped pattern (Figure 3E).

### MCI-view displays micro-domains with clustered single-molecule chromatin complexes

The proposed model of “transcription factories” that RNA polymerases are immobilized and concentrated within discrete foci in nuclei for multiple genes transcription and regulation (Cook, 1999). Data from imaging to genomics offer the evidence for RNAPII foci and RNAPII mediated promoter-enhancer clusters, i.e., “transcription factories”, participate in regulation of gene expression (Figure 4A) (Chen et al., 2016; Li et al., 2012; Wang et al., 2020). However, the corresponding genomic coordinates of individual RNAPII foci has yet to be studied, and RNAPII clusters were accumulated from bulk cells, which cannot reveal the precise composition of transcription factories (Figure 4B).

**Figure 4.**
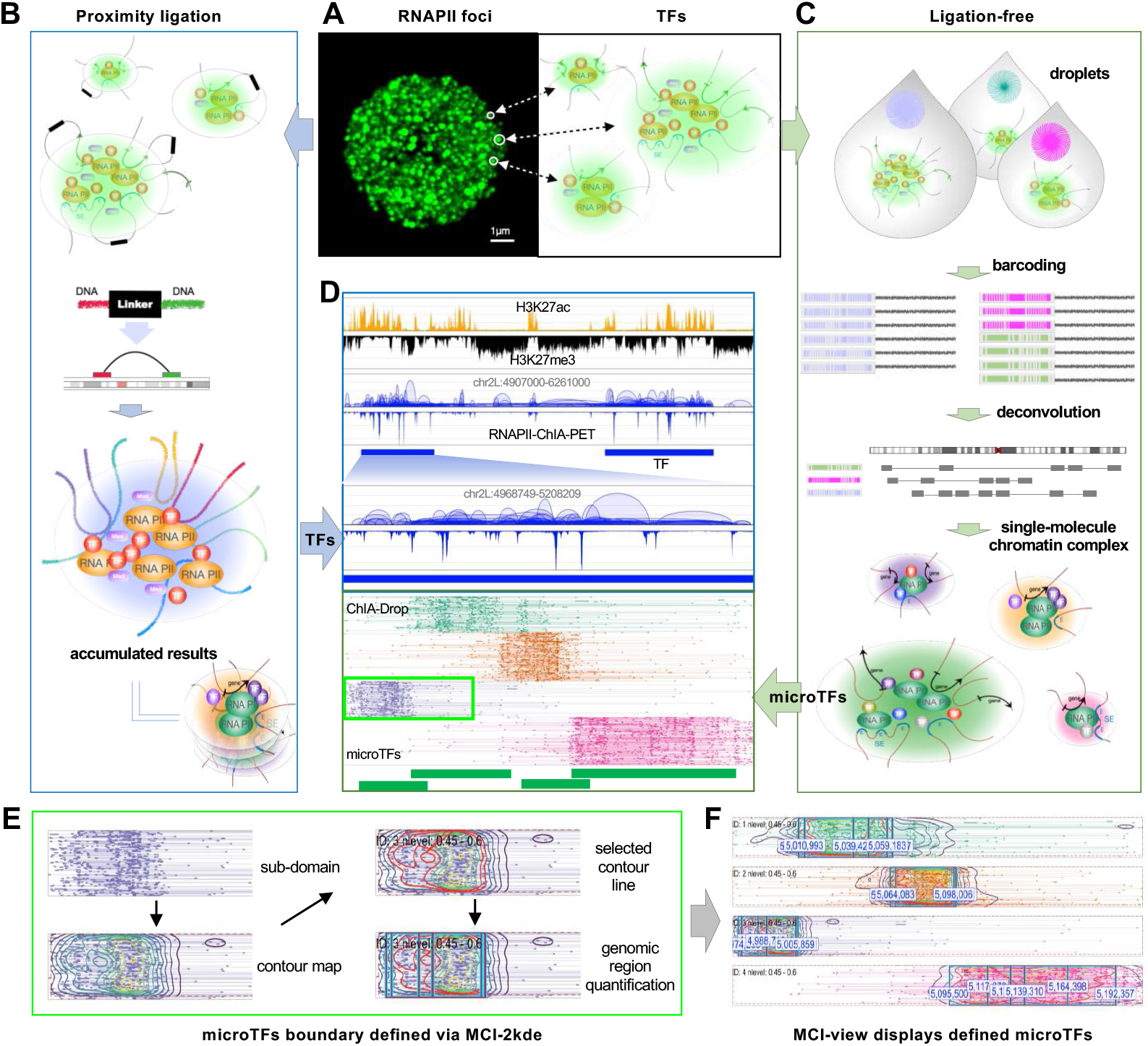
Definition of microTFs boundary by MCI-2kde. (A) RNAPII immunostaining results show different size of RNAPII foci, supporting the concept of transcription factories. (B) Proximity ligation based RNAPII ChIA-PET data demonstrated the different composition of transcription factories via high-throughput genomic sequencing strategies. (C) Ligation-free based RNAPII ChIA-Drop data disclosed the composition of transcription factories with single-molecule precision via microfluidic system to barcode individual chromatin complexes, that is all fragments of one chromatin complex within a droplet will be linked with a unique barcode. By deconvolution, the fragments that shared same barcodes would be classified to same chromatin complex, showed as linked-fragments in line. (D) BASIC Browser displays pairwise contacts of RNAPII ChIA-PET data and MCI-view displays multi-way contacts of RNAPII ChIA-Drop. RNAPII ChIA-PET data shows interaction loops and RNAPII coverage density, ChIP-seq data of H3K27ac and H3K27me3 shows the active and inactive regions, the blue bar presents transcription factories (TFs) that were defined from RNAPII ChIA-PET data. Each transcription factory is composed by several micro transcription factories (microTFs, green bar) detected from RNAPII ChIA-Drop results. (E) Definition of microTFs boundary via MCI-2kde. Density contour map was constructed by fragment distribution probability in each individual clusters shown in Fragment-view, then demonstrated the innermost contour line that covered at least 60% data points, and finally quantified their genomic coordinates by the extremums along the horizontal direction. (F) MCI-view displays microTFs boundary defined by MCI-2kde. The minimal coordinate and maximal coordinate of the square represents the boundary of the corresponded microTFs.

The advanced ligation-free chromosome conformation capture methods RNAPII ChIA-Drop could decompose the transcription factories into single-molecule precision (Figures 4C and 4D). Applying MCI-view to display the single-molecule chromatin complexes from RNAPII ChIA-Drop data, we found that the RNAPII mediated chromatin interaction domains (RAIDs) can be organized by several different micro-domains, named micro-Transcription factories (microTFs) (Figures 1G, 1H, 4D and S3A-C). This observation indicates that the micro-domains organized by a certain similarity from single-molecule chromatin contacts could potentially offer a new pathway for elucidating gene regulation (Figure S3C). We then enter the image of microTFs from MCI-view to LabelMe, a database and web-based tool for image annotation (Russell et al., 2008), for recognizing the objects of microTFs and labeling it along the boundary of microTFs. Based on LabelMe, we can obtain the ground truth boundary of microTFs by manually importing images one by one.

### MCI-2kde automatically determines microTF boundaries via density estimation

To automatically quantify the boundary of microTFs, we constructed contour density map on the microTF clusters from MCI-view by performing a two-dimensional kernel density estimation algorithm (2D KDE) (Florek and Hauser, 2010). If the points of the contour density map show the same value, they will be marked as the same contour line. Based on the density contour map of each microTF, we determined the boundary of microTFs by selecting the contour line with concentrated coverage of the fragments of chromatin complexes in Fragment-view (Figures 4E and 4F; Methods).

Using this method, we identified 578 microTFs using RNAPII ChIA-Drop data through fixed genomic regions from the previously defined 126 RAIDs with accumulated RNAPII ChIA-PET data. When we compared the boundaries of microTFs between the identification of density contour map via MCI-2kde program and that of LabelMe upon the image of microTFs from MCI-view directly, we found the average precision for 50% intersection over union (IoU0.50) is 78% when contour line setting with 60% coverage of the chromatin complexes. These indicate the approaches allow us to determine the microTFs.

In summary, we developed a framework for helping scientists to automatically define the boundary of microTFs. This framework was integrated in MCIBox and can be easily extended to the detection of other types of micro-domains generated from 3D genome mapping technologies for exploring new aspects of higher-order chromatin structure.

### Characterization of micro-domains from single-molecule chromatin complexes

The genomic feature of microTFs shown in the density plot with a peak of concentrated gene number at 6 in individual microTFs, with genomic size at a peak of 30-kb length (Figure 5A). The previously defined RAIDs by pairwise data were composed with microTFs by single-molecule chromatin complexes. We found that the genes from different phases of a cell cycle are distributed at different microTFs within a RAID, for example, the subG1 phase associated *CG17209* is located in the microTF on the left side and the G1 phase related *Myb* is located in another microTF on the right side (Figure 5B) (Björklund et al., 2006). Additionally, the overG2 phase associated *scra, sax* and G1 phase associated *tor* are all located in individual microTFs that within in a RAID (Figure 5C) (Björklund et al., 2006). Overall, the phase-specific genes of a cell-cycle could be identified in different microTFs via single-molecule chromatin contacts while before being regarded as in a same RAID by pairwise chromatin interactions (Figure 5D).

**Figure 5.**
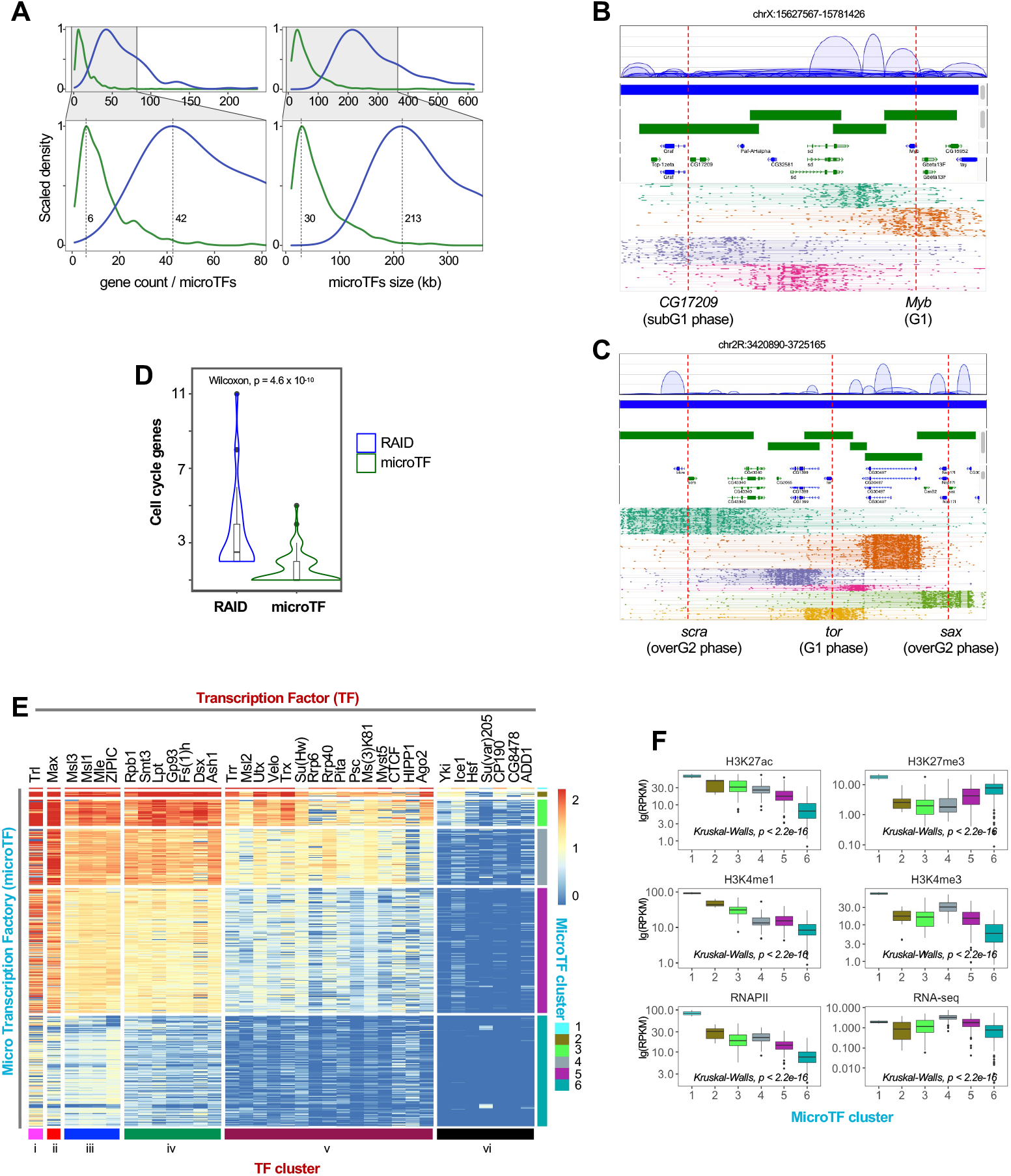
Characterization of microTFs from RNAPII ChIA-Drop data. (A) Gene count distribution. The scaled density of gene count for the defined 578 of microTFs are plotted, with a peak at 6 of genes for microTFs (green) and 42 of genes for RAIDs (blue), and the scaled density of microTFs and RAIDs size were plotted, with a peak at 30 kb and 213 kb respectively. (B-C) RAIDs (blue bar) present multiplex chromatin interactions from RNAPII ChIA-PET data are composed with several microTFs when display with RNAPII ChIA-Drop data (green bar). Panel B shows a RAID that was composed by four microTFs, left one contained cell cycle subG1 phase associated gene *CG17209*, right one contained cell cycle G1 phase related gene *Myb*. Panel C shows a RAID that was composed by six microTFs, left one contained cell cycle overG2 phase associated gene *scra*, middle one contained cell cycle G1 phase associated gene *tor*, right one contained cell cycle overG2 phase related gene *sax*. (D) The violin plots of cell-cycle gene counts for RAID and microTF was shown. The boxplot with lines as medians and whiskers at ±1.5 interquartile range; Two-sided Wilcoxon test, p value was shown. (E) We use 35 transcription factor and histone marker ChIP-seq datasets in *Drosophila* S2 cells from the modENCODE project to cluster the 578 microTFs into 6 groups by Hierarchical Clustering on Principal Components (HCPC) clustering. *y*-axis shows microTFs and *x*-axis presents transcription factors. Bar with color indicates different clusters. (F) Epigenetic feature for microTFs. The boxplot of ChIP-seq data for H3K27ac (active marker), H3K4me1(enhancer marker), H3K27me3 (inactive marker), H3K4me3 (promoter marker), RNAPII (promoter and enhancer marker) and RNA-seq data for gene expression were plotted respectively. The boxplot with lines as medians and whiskers at ±1.5 interquartile range; Two-sided Kruskal-Walls test, p value was shown.

To infer genome-wide the characteristics of microTFs, we used 35 transcription factor and histone marker ChIP-seq datasets in *Drosophila* S2 cells from the modENCODE project (Figure 5E; Methods). These 35 transcription factors are functionally clustered into different types (TF cluster in x-axis; Figure 5E), such as: (iii) Dosage compensation related factors including Msl1, Msl3 and Mle. (iv) Transcription associated factors including Rpb1, Ash1, Dsx, Smt3, Fs(1)h and Lpt. (v) Chromosome organization related factors including Rrp40, Ago2, Psc, Trr, Ms(3)K81 and CTCF. (vi) Chromosome organization regulation associated factors including CP190, Ice1 and Yki, etc. Then we investigated the association between these protein factors and the 578 microTFs by Hierarchical Clustering on Principal Components (HCPC) clustering (Figure 5E; Methods). We found these 6 clusters of microTFs are involved in different types of transcription factors. For instance, Cluster-3 and Cluster-4 consisted of 47 and 100 microTFs that were bound by most of the 35 transcription factors with different density respectively; Cluster-5 included 223 microTFs mostly bound by factors with transcription associated functions and dosage compensation related functions; and Cluster-6 with 197 microTFs showed mostly week binding by most of the factors.

To further characterize the epigenomic feature and transcriptional activity of the 6 clusters of microTFs from RNAPII ChIA-Drop data, we aligned them with histone ChIP-seq and RNA-seq data. Overall, we observed that the genes and transcription activity had high variability in the 6 clusters of microTFs (Figure 5F). As expected, the Cluster-6 microTFs exhibited low signals of H3K27ac (active histone marker), H3K4me1 (enhancer marker), H3K4me3 (promoter marker), RNAPII (transcription marker) and RNA-seq, as well as high signal of inactive histone marker H3K27me3 (Figure 5F). By contrast, Cluster-1, Cluster-2, and Cluster-3 microTFs showed strong signals of H3K27ac, H3K4me1, H3K4me3, RNAPII and RNA-seq. This observation suggests that each cluster of microTFs is involved in different types of transcription activity and histone modification.

Taken together, these results demonstrate the distinct ability and advantages of MCIBox to conveniently and effectively browse 3D genome features for the single-molecule chromatin complexes and identify the boundaries of microTFs automatically. The boundary determination of microTFs allows characterization of cell-cycle-specific or cell-type-specific gene regulation by multiplex chromatin interactions with single-molecule precision. MCIBox is an invaluable tool in making biological discoveries from many aspects for multi-way contacts data.

## DISCUSSION

In this work, we developed MCIBox for visualizing multi-way chromatin interactions and automatically identifying micro-domain boundaries. MCIBox allows for comprehensive browsing of the single-molecule multi-way chromatin interaction data generated by the cutting-edge ligation-free 3D genome mapping technologies GAM, SPRITE and ChIA-Drop. MCIBox includes an efficient MCI-view module for displaying 3D genome features and chromatin structures. MCI-view can unveil microscale structure in 3D visual panel of Cluster-view and Fragment-view, and can also facilitate exploration of the extrusion model of chromatin organization by CTCF motif-based for chromatin organization pattern view and promoter-based for transcription pattern view. Additionally, we developed frameworks MCI-2kde in MCIBox for automatically definition of the boundary of micro-domains, revealing genomic and epigenomic features of microTFs in strong association with gene regulation.

MCI-view allows for the application of a selected clustering method (hierarchical clustering, gaumix, hk-means etc.) with, or without a dimensionality reduction algorithm (UMAP, TSNE etc.) in the preceding clustering, in order to display the interactive data of multi-fragment from the ligation-free based new generation 3D genome techniques (ChIA-Drop, SPRITE or GAM) in forms of multi-type of tracks (Cluster-view, Fragment-view etc.) for regions of interest in a genome browser.

Currently, there exist more than twenty approaches for the boundary definition for chromatin structures such as TADs, yet most of them are based on profiles calculated from 2D heatmaps or epigenomic information, such as linear scores, statistical models of the interaction distributions, clustering and network features (Zufferey et al., 2018). The TAD boundaries that called by statistics often show some shifts as compared that of observation by browser directly.

To overcome these challenges, we adopted the strategy of geometric topographic map in the MCI-2kde module, where we drew a density contour map by 2D KDE in individual sub-cluster from Fragment-view and successively derived the boundary of a microTF through contour line selection. From this density contour map, we obtained 578 microTFs automatically.

As comparison of IoU region of microTF among the boundary identified approaches LabelMe and MCI-2kde, we found the percentage of their mean IoU are around 69%. Thus, the users could according to their requirement to adopt the method for boundary determination. For MCI-2kde module, the default cutoff of contour line is 60% to obtain the corresponded area covered chromatin complexes, we can increase the cutoff value by compromised for covered more chromatin complexes while with sparse distribution.

Finally, MCIBox can potentially be extended to analyze single-cell assays for higher-order chromatin structures. MCIBox, in its current form, is a convenient tool kit for single-molecule chromatin complexes analysis, which not only systemically offers a visual interface for exploring the pattern of chromatin organization, transcription, and regulation in single-molecule level, but also provides a platform for characterizing the micro-domains detected automatically from the clustered multi-way chromatin contacts. MCIBox is capable of potentially extending to distinguish the chromatin organization activity of cell-cycle-specificity and even cell-type-specificity using single-molecule chromatin contacts, to identify the chromatin extrusion model and super-enhancers regulation model.

## ACKNOWLEDGMENTS

This work is supported by grants from National Natural Science Foundation of China (32170644) and Shenzhen Innovation Committee of Science and Technology (ZDSYS20200811144002008). M. Fullwood is supported by the National Research Foundation Singapore and the Singapore Ministry of Education under its Research Centres of Excellence initiative and by a Ministry of Education Tier II grant awarded to M.J.F (T2EP30120-0020). D. Plewczynski is co-supported by Warsaw University of Technology within the Excellence Initiative-Research University (IDUB) programme, co-supported by Polish National Science Centre (2019/35/O/ST6/02484 and 2020/37/B/NZ2/03757) and EU-funded the Marie Sklodowska-Curie action (MSCA) Innovative Training Network. The authors are grateful to Y. Ruan for suggestions on manuscript organization, D. Capurso for suggestions on the improvement of MCI-view functions. The authors thank M, Kim for editing and polishing this manuscript and F. Zeng, X. Chen, H. Chen, Q. Zhu, Z. Zhang, J. Gao, M. Zhang, L. Feng, J. Bei, W. Jia, F. Li, L. Yang, E. Aiden, W. Lv, L. Liu, X. Shen and Y. Zeng for helpful discussion on this project.

## AUTHOR CONTRIBUTIONS

Conceptualization: S.Z.T., W.C. and M.Z.; Methodology: S.Z.T., G.L., Z.D., M.F., W.C., and M.Z.; Software: S.Z.T.; Investigation: S.Z.T. and M.Z.; Writing—Original Draft: S.Z.T. and M.Z.; Writing—Review and Editing: S.Z.T., G.L., M.F., D.P., J.Z., W.C. and M.Z.; MCI-view testing: D.N., Y.X., Y.Y., G.H., W.W., Z.W. and Y.L. with the assistance of S.Z.T. and M.Z. MCI-2kde testing: K.J., P.Y., and Y.L. under the direction of S.Z.T. Funding Acquisition: M.Z. All co-authors read and approved the manuscript.

## METHODS

### Data availability

The publicly available datasets used in this study are as follow: ChIA-Drop data from Zheng et al (Zheng et al., 2019) (GEO: GSE109355); SPRITE data from Quinodoz et al.(Quinodoz et al., 2018) (GEO: GSE114242); GAM data from Beagrie et al.(Beagrie et al., 2017) (GEO: GSE64881); and the available datasets for ChIP-seq data of transcription factors are shown in Table S1.

### Code availability

The source code of **MCIBox** can be accessed at https://github.com/tianzhongyuan/MCIBox. This GitHub repository includes 3 directories: **MCI-view** for visualization and micro transcription factories (microTFs) defining code by density contour map; **MCI-2kde** is the machine learning module for microTF boundary identification. The detailed code dependency and operation manual can be found in MCIBox GitHub.

### Data processing and analysis details

#### MCIBox includes MCI-view, and MCI-2kde

MCIBox toolset realizes two main functions. The function for single-molecule or single-cell data clustering visualization, and the function to define micro-domains such as micro transcription factories (microTFs) by density contour map (MCI-2kde), are realized in MCI-view module. MCI-view is coding in the framework of R Shiny Server (https://shiny.rstudio.com), which is convenience to build an interactive web application directly from R script. Without special description, R packages used in this work are from CRAN package repository (https://cran.r-project.org), where we can find source codes and description documents.

#### Data preparation for MCI-view

Currently, MCI-view main data are sourced from “ligation-free” new generation 3D genome techniques: ChIA-Drop, SPRITE and GAM (Figure 1B), which capable to capture multiplex interactive fragments from a shared chromatin complex (Figure 1C). Because the data format of these techniques out of their pipelines are different, we programed interface tools to transform these data into a uniform RGN format that MCI-view required. A RGN file actually is a fragment list, in which each data line containing a fragment with four basic columns (chromosome, start, end, and barcode) representing fragment genomic region coordinates and with a complex identity (complex barcode ID). Interface tools transform each source data into a set of RGN files composed by one chromosome per file for easier accessibility. All RGN files used for MCI-view are stored in RDS format, a binary format of R objects with a quick data loading speed and smaller storage space.

#### MCI-view workflow and visualization tracks

MCI-view is the visualization module of MCIBox toolset, and includes many types of tracks to show data profiling (Figures 1 and S1). It includes four main steps: (i) **Data Input Step**: At the very beginning, MCI-view adopts data of an interested genomic region from the according chromosome RGN file of the source library selected. (ii) **Data Format Step**: Then the input data is integrated as barcoded linked-reads and binned to matrix, formed as rows are barcodes representing complexes, columns are (original or binned) genomic region (Figures 1D and 1E). There are also three types of displays that based on data accumulation of bin-bin or bins, could be generated in this step: 2D heatmap, 2D loop or 1D coverage (Figures S1A-C). (iii) **Data Clustering Step**: MCI-view performs clustering function in either of the two strategies. The first strategy is clustering high-dimensional data (Figure 1F), which directly runs hierarchical clustering upon the rows of the high-dimensional (HD) matrix constructed from the former step. HD Cluster-view and HD Fragment-view are for viewing clustered high-dimensional data (Figures S1D and S1E). The second strategy is clustering low-dimensional data (LD, Figure 1G), that runs clustering after dimensionality reduction by assembling seven dimensionality reduction algorithms and seven clustering algorithms comprehensively. In the second strategy, MCI-view supplies selection of the combination of two types of algorithms (e.g. UMAP plus hk-means), following the principle of obtaining proper separated groups with unique colors denoted by clusters in the LD-clustering scatter plot (Figure S1F). LD-clustering scatter plot, LD Cluster-view and LD Fragment-view are tracks for clustering low-dimensional data visualization (Figures S1F-H). (iv) **Data Filtering Step**: MCI-view supplies functions for users to select data by genomic locations, such as: “Target loci” for DNA-FISH Probe-based loci filtering, “Chromatin organization pattern” for CTCF Motif-based loci filtering, “Transcription pattern” for Promoter-based loci filtering and “Transcription regulation pattern” for Super-enhancers loci filtering etc. Users can also filter data by selecting out un-interested clusters to achieve a clean “Mask-based” data view (Figure S1L).

#### MCI-view implemented clustering algorithms

We totally integrated seven clustering methods for visualization in MCI-view: Hierarchical clustering, K-means clustering, Density-Based Spatial Clustering of Applications with Noise (DBSCAN), Hierarchical Clustering on Principal Components (HCPC), Hierarchical k-means clustering (hk-means), Hierarchical DBSCAN (HDBSCAN), Gaussian Mixture Model clustering (GMM), and Fanny (a fuzzy clustering method) (Sarker, 2021).

In the area unsupervised machine learning, two important concepts need to be declared in advance: similarity and dissimilarity. They are essential in classification and clustering areas. A similarity measures how close two distributions are, while a dissimilarity denotes how different of the two vice versa. Commonly, a distance is used to quantify the similarity, such as Euclidean distance, Manhattan distance etc. A distance-based clustering algorithms tends to arrange similar data points into the same clusters, while dissimilar data points are arranged into different clusters (Shirkhorshidi et al., 2015).

Hierarchical clustering (Murtagh and Contreras, 2012) is a kind of structed connectivity-based clustering algorithm, which is based on the idea that more closely related objects should have more smaller core-distance. Initially, each object is assigned to an individual cluster; then continuously the closed clusters are joined; until only one cluster is obtained finally. At each iteration, clusters distances are refined by the Lance– Williams dissimilarity update formula based on the clustering method being selected. The formula is

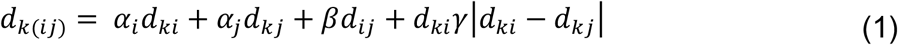

where *d_ij_* is the distance (or dissimilarity) between cluster *i* and *j, d*_*k*(*ij*)_ is the distance between cluster *k* and the new formed cluster by joining cluster *i* and *j, α_i_, α_j_, β* and *γ* are parameters that based on the applied clustering method. In this work, we implement a R package heatmap to perform a bottom-up hierarchical cluster analysis by using a dissimilarity matrix dataset to build a dendrogram.

K-means clustering is a method of vector quantization, originally from signal processing, which attempts to class a large number of objects into clusters with similar data count, in which each object belongs to the cluster with the nearest centroid k-means procedure (Wu et al., 2008). K-means clustering should include followed steps: (i) Set a *k*, which is the cluster number of the objects prepared to be classed, and randomly select *k* objects as the initial centroids. (ii) Calculate the distances between each object to all centroids, and assign every object to its closest centroid. (iii) Assign a new centroid for each of the *k* clusters by calculating mean values of two dimension respectively, so called k-means. (iv) Iterate steps (ii)-(iii), until the clusters members and centroids are stable. In a math description, given a set of data, (*x*_1_, *x*_2_,…, *x_k_*), each of these points is a real vector in *d* dimensions, and k-means is to cluster the *n* points into *k* (≤ *n*) classes, where S = (*S*_1_, *S*_2_,…, *S_k_*) to yield the minimum variances in each cluster. Generally, with the Euclidean distance, the goal is to achieve

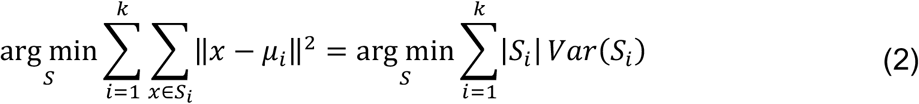

where *μ_i_* is the mean of the points in *S_i_*. k-means is a greedy algorithm to minimize the squared deviation of the data points in the same cluster:

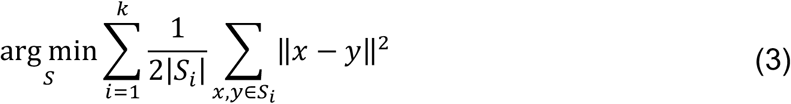

This work uses kmeans function of stats R package to realize the K-means clustering algorithm.

DBSCAN method is a density-based clustering algorithm that can be used to discovery clusters problem with irregular shapes, even in multi-dimensions. It uses the high-density connectivity of the clusters and seeks a high-density area that is separated by a low-density area. Given a set *P* of *n* points in a d-dimensional space 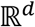, the aim is to class these points into clusters. DBSCAN defined “density” by resorting to two parameters, *ϵ* and *MinPts*. Let *B*(*p, ϵ*) be the d-dimensional spatial ball centered at point *p* with radius *ϵ*, where the distance metric is Euclidean distance. *B*(*p, ϵ*) is “dense” if it covers at least *MinPts* points of *P*. DBSCAN calls clusters following the following strategy: If *B*(*p, ϵ*) is “dense”, all the points in *B*(*p, ϵ*) should be added to the same cluster as point *p*. This creates a “chained effect”: whenever a new point 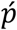 with a dense 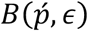 is added to the cluster of *p*, all the points in 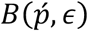 should also join the same cluster. The cluster of *p* continues to grow in this manner to the effect’s fullest extent(Schubert et al., 2017). This work uses dbscan R package to realize the DBSCAN clustering algorithm.

GMM clustering is a clustering application of gaussian mixture models (GMM), which can be seen as inducing a GMM probability concept to a k-means clustering. The GMM assumes that the data of each cluster subject to the gaussian distribution, and the overall distribution of the data is the result of the superposition of each cluster’s gaussian distribution. The use of the expectation-maximization (EM) algorithm to fit finite gaussian mixtures is currently one of the most widely used methods (Lucero-Álvarez et al., 2021). Theoretically via increasing model number, GMM could be used to approximate any probability distribution. The definition of GMM clustering is

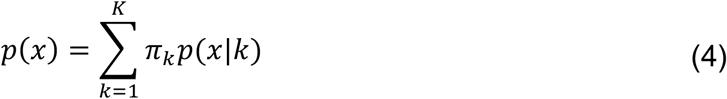

Where K is the number of models. *π_k_*is the weight of the *kth* gaussian distribution. *p*(*x*|*k*) is the *k^th^* gaussian probability density function with mean *μ_k_* and variance *σ_k_*. Now the aim of GMM calculation is to find the best *π_k_, μ_k_* and *σ_k_*, and this problem is solved by maximum likelihood estimation using EM algorithm. The iterative process of the EM algorithm is: The first step is to assume that we know the parameters of each gaussian model (one can be initialized, or based on the results of the previous iteration), to estimate the weight of each gaussian model; The second step is to determine the parameters of the Gaussian model based on the estimated weights. Repeat these two steps until the fluctuation is very small, approximately reaching the extreme value. This work uses Mclust function of R package mclust to realize the GMM clustering algorithm.

HCPC clustering executes clustering and uses the complementary between clustering and principal component methods to better highlight the main feature of the data set. This function allows to perform hierarchical clustering and partitioning on the principal components of several methods, to choose the number of clusters, to visualize the tree, the partition and the principal components in a convenient way. Finally, it provides result of the clusters (Husson et al., 2010). HCPC clustering is realized by the FactoMineR package of R.

Hk-means clustering using a hybrid approach of a combination from the hierarchical clustering and the k-means clustering. Because k-means clustering is very sensitive to this initial random selection of cluster centers, Hk-means is created for improving k-means results. Hk-means algorithm clustering data via three steps. (i) Compute hierarchical clustering and cut the tree into k-clusters. (ii) Compute the mean center of each cluster. (iii) Compute k-means by using the set of cluster centers defined in step (ii) as the initial cluster centers. In this work, the hkmean function is from factoextra package of R language.

HDBSCAN extends DBSCAN clustering method by converting it into a hierarchical clustering algorithm, and then using a technique to extract a flat clustering based in the stability of clusters. HDBSCAN clusters the data via following steps. (i) Transform the space according to the density. (ii) Build the minimum spanning tree of the distance weighted graph. (iii) Construct a cluster hierarchy of connected components. (iv) Condense the cluster hierarchy based on minimum cluster size. (v) Extract the stable clusters from the condensed tree. This work uses hdbscan function of dbscan R package to realize the HDBSCAN clustering algorithm.

Fanny is a kind of fuzzy clustering method. Unlike crisp clustering methods, fuzzy clustering often allows one object to be partitioned into more than one cluster. Fanny aims to minimize the objective function, *C*, that is to find for each object the best membership vector describing the membership coefficient of each object to each cluster.

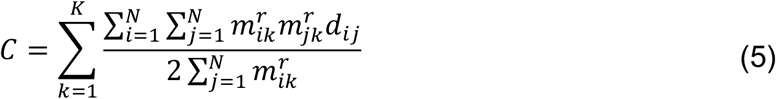

where *n* is the object number, *k* is the cluster number, *r* is membership exponent and *r* = 2 corresponding to a fuzzy c-means method, *d_ij_* is the dissimilarity between objects *i* and *j* (e.g. Euclidean or Manhattan distances). This objective function reports the nearest crisp clustering as result (Klawonn et al., 2015). In this work the fuzzy function is from cluster R package.

#### MCI-view implemented algorithms of dimensionality reduction

We assembled some dimensionality reducing algorithms in MCI-view, for the purpose of getting better clustering. they are: t-Distributed Stochastic Neighbor Embedding (TSNE), Uniform Manifold Approximation and Projection (UMAP), Principal Component Analysis (PCA), Independent Component Analysis (ICA), Multi-Dimension Scaling (MDS), AutoEncoder (AE) and Potential of Heat-diffusion for Affinity-based Trajectory Embedding (PHATE). These dimensionality reducing algorithms were arranged in front of clustering computation in the clustering on low-dimensional data strategy.

TSNE is a dimensionality reduction algorithm for non-linear data representation that yields a low-dimensional distribution from high-dimensional data. It tends to reveal local structure in high-dimensional data. TSNE is based on the stochastic neighbor embedding (SNE) model, which converting the high-dimensional Euclidean distances between data points into conditional probabilities that represent similarities. The TSNE algorithm firstly calculates a similarity measure between pairs of data points in the high-dimensional space and in the low-dimensional space, then attempts to optimize these two similarity measures using a cost function (Linderman et al., 2019). TSNE algorithm is realized by Rtsne function of Rtsne R package.

UMAP is a dimensionality reduction algorithm that can be used for general non-linear data representation. Unlike TSNE, UMAP retains more global structures with superior run-time performance. The algorithm is founded on the assumption that the data are uniformly distributed on the Riemannian manifold, of which the Riemannian metric is locally constant and the manifold is locally connected (Becht et al., 2019). Based on these assumptions, manifold with a fuzzy topology is ready for application. The embedding is found by searching a projection on the low-dimensional space owning the closest equivalent fuzzy topological structure. UMAP includes two modeling procedures: (i) building a particular weighted k-neighbor graph using the nearest neighbor descent algorithm, and (ii) computing a low-dimensional representation which can preserve desired characteristics of this graph (Xiang et al., 2021). The UMAP algorithm is realized by umap function of umapr package in R.

PCA is a popular data analysis method, which is also often used for dimensionality reduction. In the PCA dimensionality reduction process, the essence is to project the original data into a new space, which can be regard it as solving questions of the eigenvectors and eigenvalues. PCA extracts the main feature components of the data, by detecting dominant profiles and the linear combinations of the original data with maximum variance. PCA aims to find the first principal component with the largest variance and then seek the second component in the same way, which is uncorrelated with these “largest” variances. This process repeats until the new component is almost ineffective (Xiang et al., 2021). In this work, MCI-view has a function to let the user select any two dimensions to plot the low-dimensional patten. PCA for dimensionality reduction is realized by prcomp function of stats package in R.

ICA is a statistical calculation technique used to reveal the factors behind random variables, measured values, and signals. Unlike PCA preferring gaussian distributed samples, ICA aims at seeking a linear representation of non-gaussian data so that the components are statistically as independent as one can. ICA requires the source signal are of a non-gaussian distribution, and independent to each other (Choi et al., 2005). ICA for dimensionality reduction is realized by icafast function of ica package in R.

MDS is a distance-preserving manifold learning method, which can serve as a dimension reduction technique. MDS requires that the distance between the high-dimensional objects be maintained in the low-dimensional space after dimensionality reduction. (i) MDS requires to calculate a matrix of pairwise distances of a given high-dimensional data at first, (ii) then to use distance scaling to find lower-dimensional data relationship, whose pairwise distances reflect the high-dimensional distances as well as possible. The distance scaling is a nonlinear competitor of principal components, whereas classical scaling is identical to principal components (Buja et al., 2008). MDS for dimensionality reduction is realized by cmdscale function of stats package in R.

AE is a traditional dimensionality reduction algorithm, which is an artificial neural model that involves encoder and decoder model of unsupervised learning. For an AE, the encoder transforms the input high-dimensional data to lower-dimension and decoder takes the lower-dimensional data and tries to reconstruct the original high-dimensional data aiming to minimizing reconstruction error (Charte et al., 2018). AE for dimensionality reduction is realized by autoencode function of ruta package in R.

PHATE supplies a data reduction method specifically designed for visualizing high dimensional data in low dimensional spaces, using diffusion limited aggregation for generation of an artificial tree-like structure. Specially for a single-cell RNA-seq data, after PHATE calculates local affinities between cells, the affinities are used to define transitional probabilities and spread them by a Markovian diffusion framework over the data. This process causes the data dispersed and shrink onto diffusion trajectories, which are spread among several orthogonal directions identified by the eigenstates of the diffusion process. However, unlike TSNE, which conserves local structure of higher-dimensional data often at the cost of the global structure, PHATE preserving both local and global affiliations between data-points to precisely reflect the high-dimensional dataset (Moon et al., 2017). PHATE for dimensionality reduction is realized by phate function of phateR package in R.

#### Manually define microTF boundary by LabelMe

LabelMe is an image annotation tool that allows researchers to label images by hand and obtain the annotation information (Russell et al., 2008). In this work we drew a square for each micro-domain as its ground truth boundary and marked in a JSON file.

#### MCI-2kde defines microTF boundary

To automatically obtain the boundaries from micro-domains of each RAID in a Fragment-view, our initial idea was to draw a contour map according to its data distribution for each micro-domain and then select a suitable level contour line to limit the boundary. The module of MCI-2kde includes: (i) To construct contour density map for a sub-cluster (micro-domain), MCI-view only takes fragments as data points of a scatter plot despite lines. For each data point, the fragment genomic locus (coordinate of the fragment start position) and the clustered y-axis position of the complex it belongs to, are considered as two indices (*x* and *y*). (ii) MCI-view then performs two-dimensional kernel density estimation (2D KDE) over these points within a sub-cluster and returns a result of a matrix of density estimation approximation, consequently the matrix displays the results as contour maps. (iii) Finally, a contour line that encircles the majority of data points of a micro-domain is selected for the microTF boundary definition.

In this work, MCI-view employs geom_density_2d program (ggplot2 package of R) to call kde2d function perform the 2D KDE over a micro-domain fragments and outputs as a contour density map, which generates 10 levels of contours by default. Next, the innermost contour line that enclose at least 60% data points was selected. Finally, the region between the leftmost points to rightmost points of the selected contour line is squared to represent the micro-domain boundary.

#### Clustering of the matrix crossing microTFs and transcript factors (TFs)

To study functions of microTFs, the cell-cycle associated genes described previously (Björklund et al., 2006) were assigned to RAIDs and microTFs according to their coordinate respectively, then gene count in individual RAIDs and microTFs were plotted as violin plot (Figure 5D). In addition, we selected 35 ChIP-seq data of TFs from literatures (Table S1), and assigned their coverages to each of the auto-defined 578 microTFs to construct a matrix. Next, we performed Hierarchical Clustering on Principal Components (HCPC) clustering on both directions of the matrix, and achieved 6 groups of microTFs and 6 groups of transcription factors (TFs) (Figure 5E), which are displayed by heatmap in Figure 5E. Subsequently, we also used data of histone marks, RNAPII ChIP-seq and RNA-seq from literatures (Table S1) to hit microTFs, and shown as boxplots faceted by different microTF clusters in Figure 5F.

